# Analysis of Nucleotide Pools in Bacteria Using HPLC-MS in HILIC Mode

**DOI:** 10.1101/655258

**Authors:** Eva Zborníková, Zdeněk Knejzlík, Vasili Hauryliuk, Libor Krásný, Dominik Rejman

## Abstract

Nucleotides, nucleosides and their derivatives are present in cells at varying concentrations that rapidly change with the nutritional, and energetic status of the cell. Knowledge of the concentration dynamics of these molecules is instrumental for understanding their regulatory effects. Determination of these concentrations is challenging due to the inherent instability of these molecules and, despite many decades of research, the reported values differ widely. Here, we present a comprehensive and easy-to-use approach to determine intracellular concentrations of >25 target molecular species. The approach uses rapid filtration and cold acidic extraction followed by high performance liquid chromatography (HPLC) in the hydrophilic interaction liquid chromatography (HILIC) mode using zwitterionic columns coupled with UV and MS detectors. The method reliably detects and quantifies all the analytes expected to be observed in the bacterial cell and paves the way for future studies correlating their concentrations with biological effects.

## Introduction

Nucleotides and their derivatives play crucial roles in many biological processes in bacterial cells. They serve for energy storage or as building blocks for the assembly of macromolecules such as nucleic acids or polysaccharides. Furthermore, nucleotides play important regulatory roles, e. g. functioning as second messengers and/or pleiotropic regulators: cyclic AMP (cAMP) [1], cyclic diadenylate (c-di-AMP), cyclic diguanylate (c-di-GMP) [2], dinucleoside polyphosphates ApxN [3], as well as guanosine pentaphosphate (pppGpp) and tetraphosphate (ppGpp), collectively referred to as (p)ppGpp [4, 5]. Rapid increase in (p)ppGpp concentration – ‘the stringent response’ – orchestrates a survival program leading to increased virulence and antibiotic tolerance [6]. In *Escherichia coli* the levels of (p)ppGpp are controlled by two related enzymes – RelA and SpoT, the namesakes of RelA SpoT Homolog (RSH) protein family – that sense numerous stress signals such as amino acid, fatty acid, carbon source limitation *etc*. [7]. The (p)ppGpp then functions as a pleiotropic regulator binding to, and affecting the activity of many *E. coli* targets, including RNA polymerase, GTPase enzymes involved in protein synthesis and ribosome assembly as well as nucleotide biosynthesis enzymes [8-11]. In addition to (p)ppGpp, related pGpp and pGp nucleotides have been detected in cellular extracts, though their biological significance is not clear and analytic differentiation from GTP and GDP is not trivial [7].

Detection and quantitation of nucleotide pools in bacterial cells consists of three steps: (i) sample acquisition, (ii) extraction and (iii) analysis. During sample acquisition, it is advantageous to separate bacterial cells from the medium. Generally, the whole culture approach results in less sensitive analysis, since the samples are more diluted and include cultivation broth compounds, which results in high background noise. Cells-medium separation then can be done either by centrifugation or filtration. The centrifugation method has been used in metabolomic studies e.g.[12]. However, filtration is superior since it is faster and less prone to affecting the physiology of the cells, and therefore for nucleotide analysis centrifugation should be avoided since it dramatically perturbs the nucleotide pools [13].

The next step is extraction. It is done by cold or hot organic solvent (usually methanol or its mixture with water, acetonitrile or chloroform) [14, 15], acids (formic [16] or acetic [13]) or alkali [17, 18]. In the whole culture sampling approach, the extraction step is omitted, which can be beneficial for some highly labile analytes.

The third step – analysis – is the most challenging one. It was historically done on thin layer chromatography plates developed in two dimensions (2D-TLC) [16]. This method is relatively simple in terms of technical equipment but rather laborious and time-consuming. Moreover, the migration of standards and identified analytes could be misinterpreted, e.g. separation of nucleotide triphosphate from their deoxy counterparts. Finally, quantification of analytes by this method lacks precision. Alternatively, anion Exchange Liquid Chromatography (AEX) [13, 15, 19] and Ion-Pair Liquid Chromatography (IP-LC) [13, 15, 18, 20] have been widely used in nucleotide analysis for decades. Both approaches provide very good separation according to analyte charge and size and detection with UV detectors afford satisfactory quantification of many analytes based on comparison with known standards. However, all analytes have to be baseline-separated for proper quantification and co-eluting compounds that happen to absorb at the same wavelengths could lead to errors in quantification. This can be solved by coupling IP-LC to a mass spectrometry (MS) analysis [21-23]. This approach benefits from well separated analytes along with selective detection. However, using ion pair reagents (even volatile ones) decreases MS signal intensity [24] and renders the MS spectra more complex due to adduct formation. Furthermore, the need to frequently clean MS detectors when ion pair additives are used adds to the work required by this approach. Hence, hydrophilic Interaction Liquid Chromatography (HILIC) appears to be the method of choice for separation of small polar compounds. The major advantage over AEX or IP-LC is the possibility to use MS and thus obtain sensitive and selective detection outputs. AEX could be connected to an MS detector only by using electrolytic suppression [25]. The high organic content in HILIC MS simplifies electrospray evaporation and increases the MS signal intensity. Zwitterionic columns (ZIC-HILIC) represent one type of these columns, where either sulfoalkylbetaine functional groups are bonded to a silica gel support (ZIC-HILIC) or polymer support (ZIC-pHILIC), or where the oppositely charged phosphorylcholine functional groups are bonded to a silica gel support (ZIC-cHILIC) (**Fig. 1**). In all cases, strong positive and negative charges are at the exact ratio of 1:1. The permanently charged stationary phase is not affected by the pH of the mobile phase, and thus the pH of the mobile phase influences only the charge of the analytes. Separation is achieved by hydrophilic partitioning combined with weak ionic interactions. HILIC-MS is mainly used in metabolomics studies [26, 27] but recently, this technique is becoming popular in nucleotide analysis [28-30]. In most cases, UV or tandem mass spectrometry (MS/MS) quantification is used. However, only a few metabolomic studies reports describing the use of a single MS but they quantify only several selected nucleotides [31, 32].

**Figure 1.**
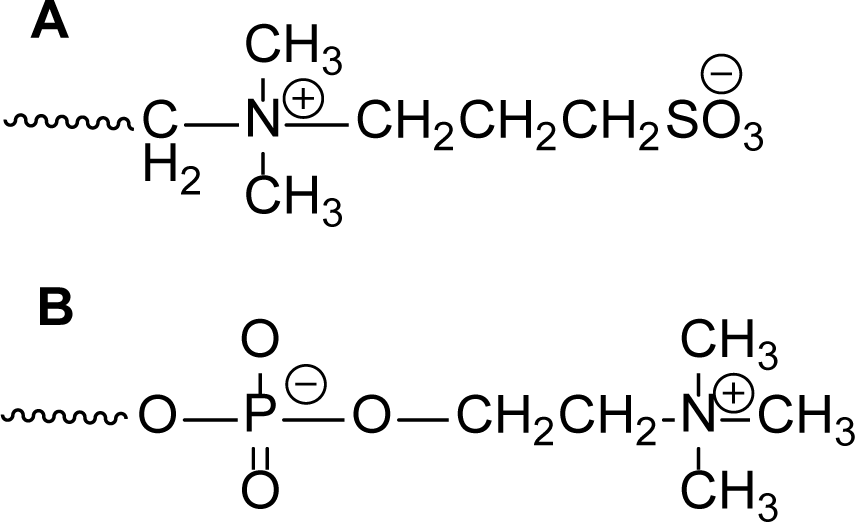
ZIC-HILIC (**A**) a ZIC-cHILIC (**B**) column chemistries.

In this study we present a comprehensive and easy-to-use optimized protocol for sensitive detection and quantification of nucleotides and their derivatives. It is based on rapid filtration of cells followed by acidic extraction and separation on high performance liquid chromatography (HPLC) using zwitterionic HILIC columns. Filtration times, six different stationary phases of HILIC columns, various gradients of the mobile phase and different additives and their concentrations and pH were extensively tested. The detection is achieved by a single MS detector – a simple yet powerful analytical tool, typically present in most laboratories. The final optimized and sensitive method yields highly reproducible data for >25 nucleotides and their derivatives, including pGpp and pGp.

## Materials and methods

### Chemicals

Mupirocin, MOPS, and tricine were purchased from Sigma-Aldrich (USA). Standards were purchased as solids from Sigma Aldrich (AMP, ADP, ATP, CMP, CDP, CTP, dCDP, GMP, GDP, cGMP, dGMP, UMP, UDP, IMP, IDP, ITP, NAD, NADH, NADP, NADPH, ^13^C-ATP) or as a aqueous solutions from Sigma Aldrich (dTTP, dCTP) or Jena Bioscience (dGTP, dITP, dATP, UTP, GTP, ppGpp and pppGpp). Unusual species were synthesized in our lab (ppGp, pGpp, pGp) [according to modified method of [33] as well as internal standard (IS) **ProG** (**Fig. 2**) [34]. HPLC grade solvents were purchased from VWR (Acetonitrile, Methanol), UPLC-MS grade solvents from Merck (Acetonitrile). Both HPLC and UPLC grade water was obtained via MilliQ filtration system (Merck) with appropriate filters.

**Figure 2.**
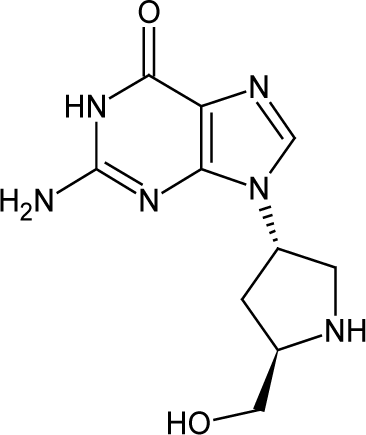
Structure of prolinol nucleoside analog **ProG** which was used as an internal standard. Calculated LogD_5_ is −4.8 and logD_7_ is −5.1 (as per Marvin calc).

### Preparation of standard solutions

All standards were diluted in UPLC grade water to the stock concentration of 1 and 10 mmol/L and stored at −20 °C. NADH and NADPH are very labile and stock solution was used only for one week, after that fresh one was prepared from powder.

Standard mixtures were prepared from stock solutions. There were two standard mixtures: (I.) unstable compounds: NADH, NADPH, ppGpp and pppGpp; and (II.) stable compounds: AMP, ADP, ATP, CMP, CDP, CTP, dCDP, dCTP, GMP, GDP, cGMP, dGMP, UMP, UDP, IMP, IDP, ITP, NAD, NADP, dTTP, dGTP, dITP, dATP, UTP, GTP and pGp and pGpp. ppGp was not used for quantification, since we were not able to separate it properly from isomeric pGpp. These two standard mixtures were prepared at 8 concentration levels: 0.5, 1, 2, 5, 15, 30, 50 and 100 µM. We have distributed lower part of calibration into more points (1, 3, 6, 10, 15 µM) for proper determination of shape of calibration curve.

Quality control (QC) mixture was prepared based on published data [16], where defined concentrations were diluted ten to twenty times to mimic on column concentration in measured samples. QC composition is listed in **Tab. S1**.

IS was added into all standard and QC solutions, as well as to all measured samples, to final concentration of 25 mM. Final composition of all standard and QC solutions was 50% acetonitrile and 50% water.

### Columns

Six tested HPLC HILIC stationary phases were: (I.) crosslinked diol (Luna HILIC diol, 100 × 4.6 mm, 5 µm, 200 Å; Phenomenex), (II.) bare silica (Luna SiOH, 100 × 4.6 mm, 5 µm, 100 Å; Phenomenex), (III.) monolithic amino (Chromolith NH2, 100 × 4.6 mm, 2 µm, 130 Å; Merck) and two zwitterionic functional groups – sulfoalkylbetaine bonded to (IV.) polymer support (ZIC-pHILIC, 100 × 4.6 mm, 5 µm; Merck) or (V.) silica gel support (ZIC-HILIC, 150 × 4.6 mm, 3.5 µm, 100 Å; 150 × 2.1 mm, 3.5 µm, 100 Å and 250 × 2.1 mm, 3.5 µm, 100 Å; Merck) and (VI.) phosphorylcholine bonded to silica support (ZIC-cHILIC, 150 × 2.1 mm, 3 µm, 100 Å; Merck) (**Fig. 1**).

### Instrumentation

Method was developed on HPLC system with binary solvent gradient LC-20AD with single quadrupole mass spectrometer LC-MS 2020 and photodiode array (PDA) detector SPD-M20A (Shimadzu) and optimized on Acquity H-Class UPLC system with hybrid quadrupole-time-of-flight mass spectrometer (qTOF) Xevo G2-XS QTof and Acquity PDA detectors (Waters). Data evaluation was performed using MassLynx software with TargetLynx application manager (Waters).

### Analytical parameters

Analysis of *E. coli* extracts were performed on ZIC-HILIC columns (150 × 4.6 mm, 150 × 2.1 mm and 250 × 2.1 mm) with flow rates 0.5 ml/min, 0.3 ml/min and 0.15 ml/min, respectively. Mobile phase A was 10 mM ammonium acetate (NH_4_Ac) with pH adjusted to 5 with acetic acid and B was 90% acetonitrile with 10% of 100 mM acidified ammonium acetate pH 5. Induced stringent response was monitored on ZIC-cHILIC columns to separate pppGpp and ppGpp and to improve their MS signal. ZIC-cHILIC column (150 x 2.1 mm) ran at flow rate 0.15 ml/min with mobile phase A = 25 mM NH_4_Ac and B = 75% acetonitrile with 25% 100 mM NH_4_Ac.

MS quantification was performed by negative electrospray ionization with following parameters: capillary voltage = 2 kV, cone voltage = 20 V, source temperature = 120 °C, desolvation temperature = 400 °C, desolvation gas flow = 400 L/h, cone gas flow = 30 L/h.

### Harvesting of bacterial cells and extraction of nucleotides

*E. coli* K12 strain was grown on lysogeny broth (LB) agar media prior the experiment at 37 °C. Bacterial cultivation and sample processing were adapted from [13]. To prepare *E. coli* inoculum, a fresh bacterial colony on LB agar media was transferred to 2 ml of MOPS medium (40 mM MOPS / 4 mM Tricine adjusted to pH 7.4 with KOH, 1 mM glucose, 9.5 mM NH_4_Cl, 0.27 mM K_2_SO_4_, 1.3 mM K_2_HPO_4_, 0.52 mM MgCl_2_, 50 mM NaCl, 10 μM FeSO_4_, 0.002 μM (NH_4_)_6_Mo_7_O_24_, 0.4 μM H_3_BO_3_, 0.03 μM CoCl_2_, 0.009 μM CuSO_4_, 0.08 μM MnCl_2_, 0.001 μM ZnSO_4_) and incubated 16 hours at 37 °C and 220 rpm. 50 ml of the MOPS medium in 250 ml Erlenmeyer cultivation flask was next inoculated to initial OD_600_ 0.005 and bacterial suspension was cultivated at 250 rpm at 37 °C and until reaching of exponential phase, which was determined by OD_600_ time dependence in the interval of 0.05 to 0.7. To induce the RelA-mediated stringent response the cells were challenged by 20 or 150 μg/ml mupirocin for 5 to 35 min prior the sample collection. In dependence of type of experiment 5 to 8 ml cell suspension was quickly collected from cultivation flask which was quickly vacuum filtered thorough 25 mm 0.45 μm cellulose acetate filter. Membrane with detained bacteria was immediately transferred to 1 ml ice-cold 1M acetic acid in 1.5 ml microtube and frozen in liquid nitrogen. Samples exceeding the time of both filtration and freezing more than 15 s were discarded. For each point of analysis three to five parallel samples were collected to estimate precision of the process extraction. For recovery estimation 1 M acetic acid was spiked by 5 μM ^13^C_10_-ATP as internal standard. Samples were next slowly thawed on ice and after completing thawing they were next incubated for 30 min on ice with short vortexing in 5 min intervals. Crude bacterial lysate was separated from cellulose acetate filter by short centrifugation (4,000 x g, 30 s) through pinhole at bottom of 1.5 ml microtube assembled with 2 ml collection tube. Lysate was quickly frozen in liquid nitrogen, lyophilized and material was then resuspended in 200 μl of ice-cold deionized water and 30 min incubated on ice. Insoluble material was removed by 20 min centrifugation at 22,000 x g at 4 °C. Clear aqueous bacterial extract was collected and stored at −80 °C to further analysis. For each point we have prepared three to five parallels to estimate precision. Extraction recovery was determined by adding fully isotopically labeled ATP-^13^C_10_ into extraction media of 1 M acetic acid to final (i.e. in sample) concentration of 25 mmol/L.

### Cell volume calculation

Based on the published data [13, 35] we approximated cell number of *E. coli* exponentially growing in liquid LB media at OD_600_ = 0.45 to 3.9 × 10^8^ and cell volume to 1 × 10^-15^ l.

### Validation parameters

Method was validated using ZIC-cHILIC column, 150 × 2.1 mm and ZIC-HILIC column 250 × 2.1 mm according to the guidelines defined in U.S. Pharmacopeia [36] and advises for application in bioanalytical method validation [37].

### Selectivity

First, each standard was injected separately to estimate its RT. Next, mixtures of standards with different mobilities were prepared at four concentration levels – 2, 5, 20 and 50 µM. Mixtures 1-7 were composed of: 1) AMP, ADP, ATP, dATP, NAD; 2) GTP, GDP, GMP, dGTP, dGMP, cGMP; 3) CTP, CDP, CMP; 4) UMP, UDP, UTP, dTTP; 5) ITP, IDP, IMP, dITP; 6) pGp, pGpp, ppGpp, NADP; and 7) pppGpp, NADH, NADPH. Standards were distributed to ensure all analytes would be baseline separated and their peak purity was monitored both by MS and UV. Finally, both standard mixtures, as described in *Preparation of standard solutions*, were injected at four concentration levels - 2, 5, 20 and 50 µM, to ensure that the method is capable to properly identify every analyte at various possible concentration level.

### Linearity, Limit of Detection, Limit of Quantification

Standard solutions were prepared as described in *Preparation of standard solutions* section. Primarily each standard was analyzed separately to obtain limit of detection (LOD), limit of quantification (LOQ) and also upper limit of quantification, when detector was saturated. LOD was determined as signal to noise ratio (S/N) equals 3 and LOQ as S/N equals 10. Thus, we obtained “quantification area” for these standards. We diluted stock solutions into mixed standard solutions at eight different concentration levels, that were distributed among this area. From the working solutions calibration curve for each standard was plotted as a ratio between analyte response and IS response to IS concentration. We have used linear regression for nucleoside monophosphates and quadratic regression for nucleoside di- and triphosphates with weighting 1/x.

### Accuracy and Precision

We estimated method accuracy on the QC mixture and on standard mixtures at three concentration levels [37], that we have prepared as described in *Preparation of standard solutions*. We determined precision in the terms of repeatability both for biological samples and artificial pool mixtures. Intermediate precision was determined only on QC mixture, since there was serious concern for bacterial extract stability during long-term storage.

### Robustness

We tested the influence of temperature, composition of mobile phase (concentration of buffer and its pH), injection volume and composition of injection solvent with keeping the other parameters constant on QC sample. Each point was measured three times. Influence of mobile phase composition was examined only on ZIC-cHILIC column.

1. Temperature: We heated the column to 30 °C, 35 °C and 40 °C and compared the separation to the analyses without heating.
2. Buffer concentration: We tested three concentration levels: 20 mM, 23 mM and 25 mM.
3. Buffer pH: We compared 100 mM NH_4_Ac buffer without pH modification (pH = 6.91) with acidified buffer with acetic acid (pH = 6.65) and alkalized buffer with ammonia (pH = 7.15). Gradient was as follow: 0% A, 75% B, 25% C for 3 min, 20% A, 55% B, 25% C in 20 min, hold till 25 min. A = water, B = acetonitrile, C = 100 mM NH_4_Ac.
4. Injection volume and composition of injection solvent: We compared three injection volumes: 1 µL, 2 µL and 3 µL. For each injection volume three compositions of injection solvent were tested – 25% acetonitrile, 50% acetonitrile and 75% acetonitrile.

### Matrix effects

Matrix effects (ME) were evaluated through comparing IS response of addition into standard mixture in 50% acetonitrile at various concentration levels with IS response in bacterial growth medium and bacterial extract and calculated matrix effects as: ME (%) = B/A × 100 [38]; where A is peak area in standard solution and B is peak area of the standard spiked after extraction. We also compared the ratio of MS response to UV response of synthetized IS added after extraction with the same ratio of isotopically labeled ATP (^13^C_10_-ATP) added into extraction solution. We made this juxtaposition for standard mixtures, extracted MOPs medium and bacterial extracts.

## Results

### Method development

We started developing the method by preparing a defined mixture of nucleosides and nucleotides and a suitable internal standard (IS) that is essential for quantification of the analytes (see Materials and Methods). The most important requirement for the choice of IS is its natural absence in the sample. The IS should be stable enough and mimic ionization properties of the majority of analytes. IS should absorb in the same region as the analytes for quantity control and its retention time (RT) must differ from RTs of the other analytes. We decided to use non-isotopically labeled IS due to its manipulation ease and cost considerations. As we were focusing mainly on guanosine phosphates, we chose IS with the guanine nucleobase. After evaluation of three different compounds as IS we finally selected the synthetic nucleoside analog **ProG** (**Fig. 2**) due to its best ionization properties. We always added IS to the sample after extraction and its function was to minimalize systematic error and signal fluctuation due to random changes.

Then, we tested resolution of the nucleoside and nucleotide mixtures (including IS) under 12 mobile phase conditions (four different pH levels (3, 5, 7 and 9) combined with three different solvent additives (ammonium formate, ammonium acetate, and ammonium bicarbonate)) on six different, commonly used HILIC columns (see Materials and Methods), yielding a total of 72 combinations.

This systematic screen identified two columns and conditions for the best resolution of complex mixtures of neutral, slightly polar, and very polar compounds. These two columns were the zwitterionic ZIC-HILIC and ZIC-cHILIC columns. These two columns have opposite charges on their surfaces, and, therefore, the separation mechanisms vary. On the negatively charged ZIC-HILIC column, partitioning is aided by electrostatic repulsion (ERLIC) [39] whereas on the positively charged surface of ZIC-cHILIC, partitioning is combined with electrostatic interactions. Therefore, the retention behaviors of these two columns slightly differed. Interestingly, the elution order of the majority of the analytes remained unchanged (see **Tab S2** and **S3**). The most pronounced difference in the elution order (characterized by the mobility of the analyte) was observed for IS, which eluted with the shortest RT at the ZIC-cHILIC stationary phase and with the longest RT on ZIC-HILIC. In both cases it eluted before or after the region where all UV-absorbing species eluted. Hence, the detection of any analyte of interest was not compromised.

The mobile phase affected the resolution and peak shapes of the analytes, depending on the column used. For ZIC-HILIC, slightly acidic conditions (pH 5) improved peak shapes but also decreased the MS signal level, especially in the negative ionization mode as analytes were less charged under these conditions. This phenomenon was more significant for analytes bearing more phosphate groups. Unfortunately, ppGpp and pppGpp were only marginally resolved under these conditions.

To the contrary, the ZIC-cHILIC column was able to separate well pGpp/ppGp, ppGpp and pppGpp under the slightly acidic conditions described above. Nevertheless, pH adjustment of the mobile phase to pH 7 markedly improved their MS signal. To avoid split peaks at neutral conditions for some analytes (mainly AMP or NAD) and to increase the separation factor, higher salt concentration was used (25 mM ammonium acetate instead of 10 mM) (**Fig. 3**). Ammonium ions served as weak ion pair reagents and improved reproducibility. Nevertheless, the higher salt concentration that was used prolonged the equilibration time and also resulted in more frequent MS source cleaning. This is consistent with the literature, where it was reported that the application of volatile strong ion pair reagents in analysis of polar metabolites improves separation efficiency and peak shape [38] but causes signal suppression [28].

**Figure 3.**
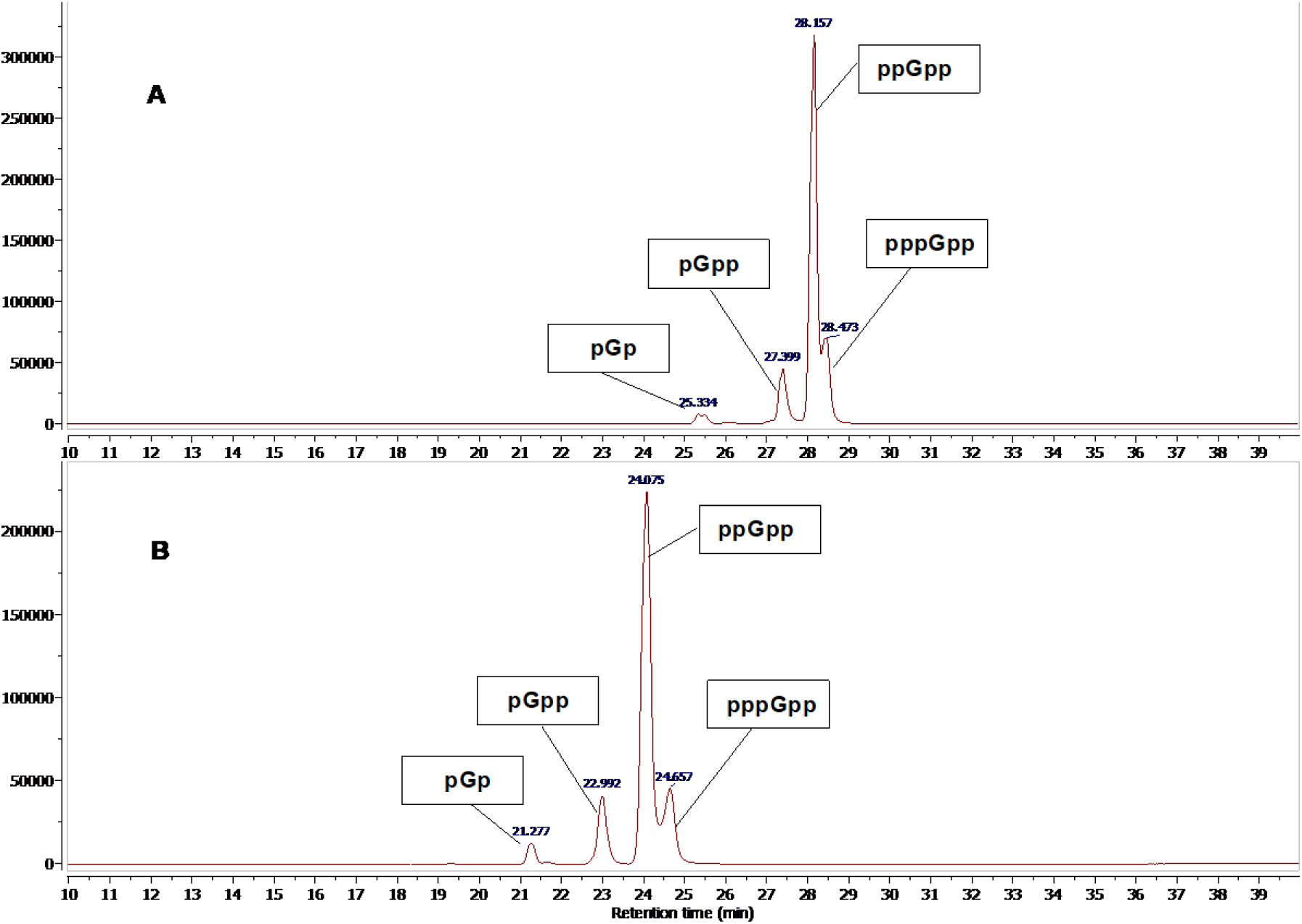
Separation of different guanosine phosphates on ZIC-cHILIC column. Column: ZIC-cHILIC, 150 × 2.1 mm, 3.5 um. Gradient condition: (**A**) 15% A, 75% B, 10% C for 3 min, 25% A, 65% B, 10% C in 20 min, 35% A, 55% B, 10% C in 25 min; (**B**) 5% A, 75% B, 20% C for 3 min, 25% A, 55% B, 20% C in 20 min, hold till 25 min. A = water, B = acetonitrile, C = 100 mM NH_4_Ac.

Using the HILIC mode of separation allowed us to avoid strong ion pair reagents. Although ammonium acetate also caused signal suppression, this suppression was more pronounced for multiply-charged analytes [24]. For singly charged analytes, such as nucleotide species, no significant signal suppression with higher ammonium acetate concentration was observed, especially in the negative ionization mode.

To conclude this part, for resolution of nucleotides and nucleosides, and especially of the stringent response alarmones (p)ppGpp, the ZIC-cHILIC column performed the best. A comparison of the performance of ZIC-HILIC and ZIC-cHILIC columns for 29 analytes is shown in **Tab. S2** and **S3**. The next part validates the ZIC-cHILIC column performance.

### Method validation

#### Selectivity

As our detection method of choice, we selected a single MS detector, a common equipment in most laboratories. Compared to UV, it is more sensitive; compared to MS/MS, it is more user-friendly, not requiring highly trained personnel.

The use of MS detection required that analytes of the same nominal mass did not co-elute (e.g. isomers of guanosine triphosphates or diphosphates, adenosine phosphates with deoxyguanosine phosphate) and that co-eluting substances did not affect their ionization. Furthermore, separation of species with the same nucleobase was also critical as phosphate loss may occur during the ionization process in the source, resulting in e.g. the concomitant presence of triphosphate, di-, and monophosphate in the spectrum (**Fig. 4**). Thus, mono- and diphosphates could be overestimated if they had the same retention time as the triphosphate. This phenomenon is even more pronounced with highly phosphorylated species as (p)ppGpp, where the signal of the species without one phosphate could be even stronger than the parent ion. In the positive ionization mode (ESI+), these effects are less significant.

**Figure 4.**
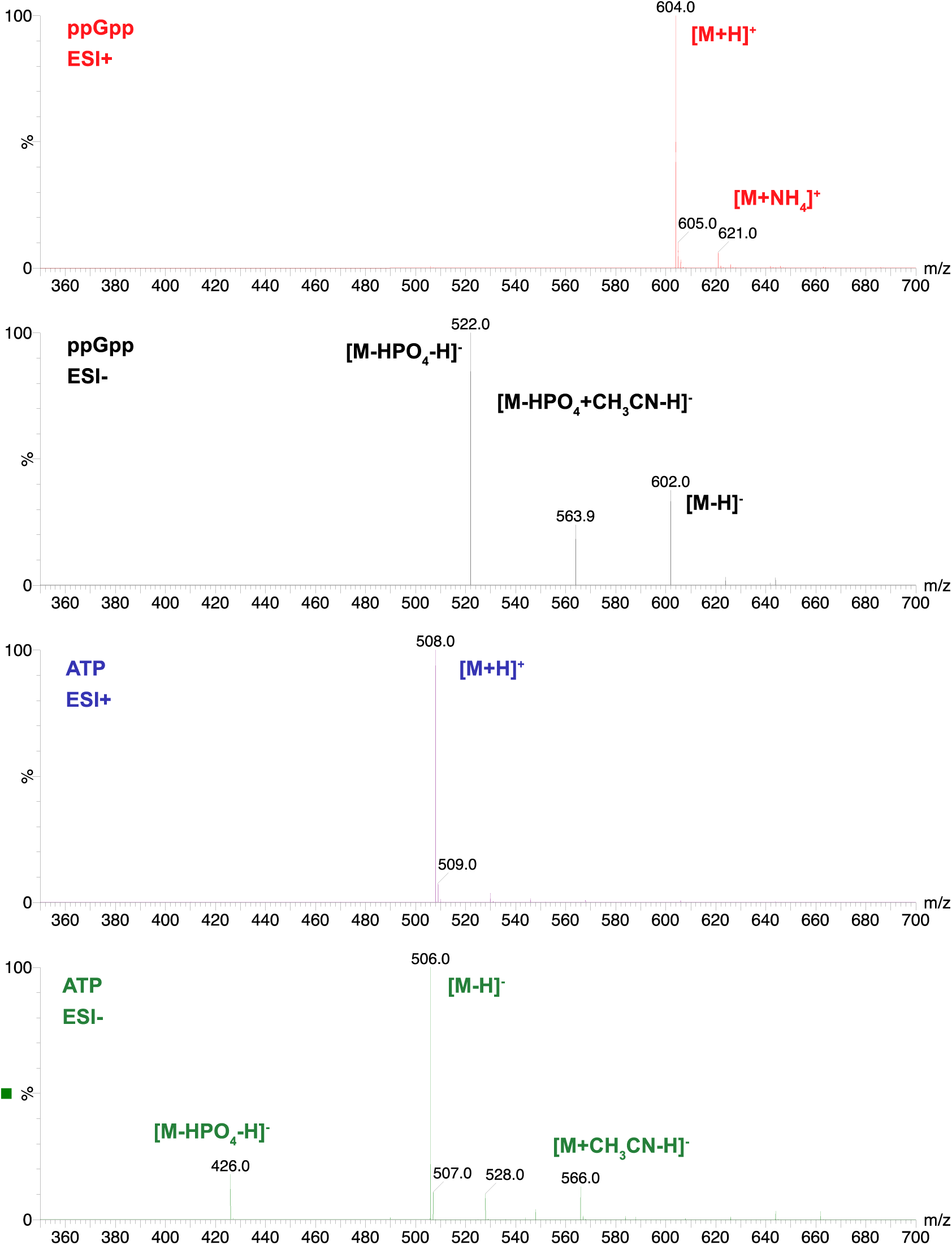
Ionization pattern of ppGpp and ATP in positive and negative ionization mode. In source phosphate loss increase with higher phosphate groups in molecule (ATP *vs*. ppGpp) and with applied ionization mode, where ESI+ induce phosphate loss in much less extent. Electrospray ionization with following parameters was used: capillary voltage = 2 kV, cone voltage = 20 V, source temperature = 120 °C, desolvation temperature = 400 °C, desolvation gas flow = 400 L/h, cone gas flow = 30 L/h.

**Tab. S4** shows the selectivity performance for selected pairs of analytes mentioned above. We baseline-separated almost all of the tested species except for ppGp and pGpp that eluted at the same time. In the case of ppGpp and pppGpp, a minor overlap was observed after ≈100 injections, due to the column aging.

#### Limit of Detection, Limit of Quantification, and Linearity

Limit of detection (LOD) in solution drifted from 0.2 µM to 2 µM. This corresponded to 0.4 to 4 pmol on the column. Limit of quantitation (LOQ) was approx. 2-5 times higher than LOD. Values for LOD, LOQ are listed in **Tab. S2 and S3.**

Linearity was validated with the calibration curve response. Not every analyte fitted the linear regression within the whole area. However, as extensively reported [40-42], it is not necessary to force the calibration data to a linear curve. In ESI-, the linear range for nucleotide di- and triphosphates is quite narrow. Therefore, using a quadratic function for these analytes covers the entire expected concentration range and provides good reproducibility. Coefficient of determination (R^2^) was in all cases > 0.98 and for the majority of analytes even > 0.99. Further discussion is in the *Calibration curve* section.

#### Precision and Accuracy

The obtained data were highly precise both in inter-day and intra-day measurements and the relative standard deviation (σ) was lower than 5%, indicating that no significant systematic error had occurred. In the case of accuracy, variance in measured value and true value (expressed as bias) increased at low concentration levels close to LOD. Nevertheless, the values were within the acceptable range according to FDA guidelines [43]. The results are listed in **Tab. S1**.

### Robustness

The influence of temperature (**Fig. 5**), buffer concentration (**Fig. 6**), buffer pH (**Fig. 7**), injection volume and composition of injection solvent were tested.

**Figure 5.**
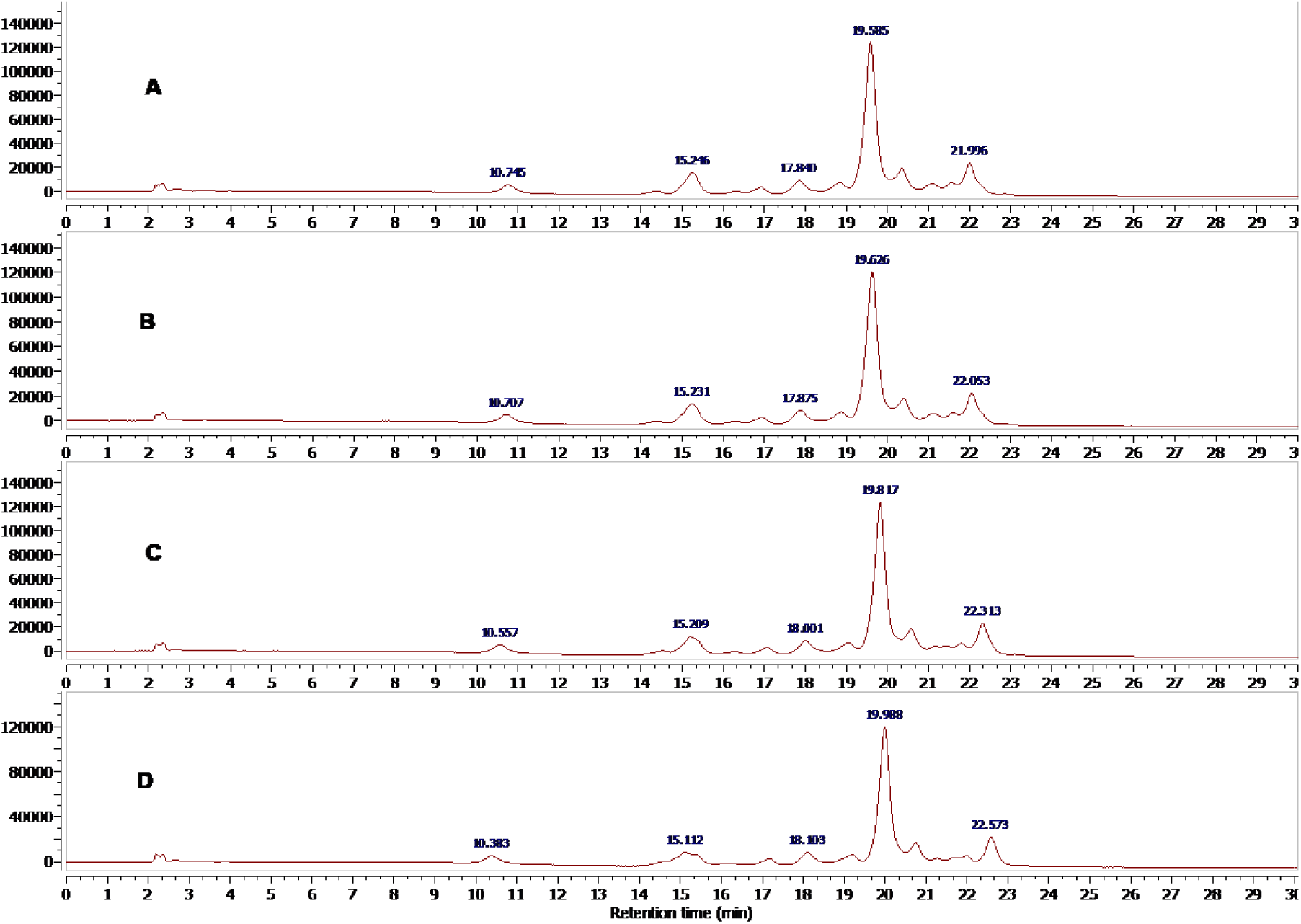
Effect of the temperature on QC sample. Column ZIC-cHILIC, 150 × 2.1 mm, 3 µm; (**A**) room temperature, (**B**) 30 °C, (**C**) 35 °C and (**D**) 40 °C.

**Figure 6.**
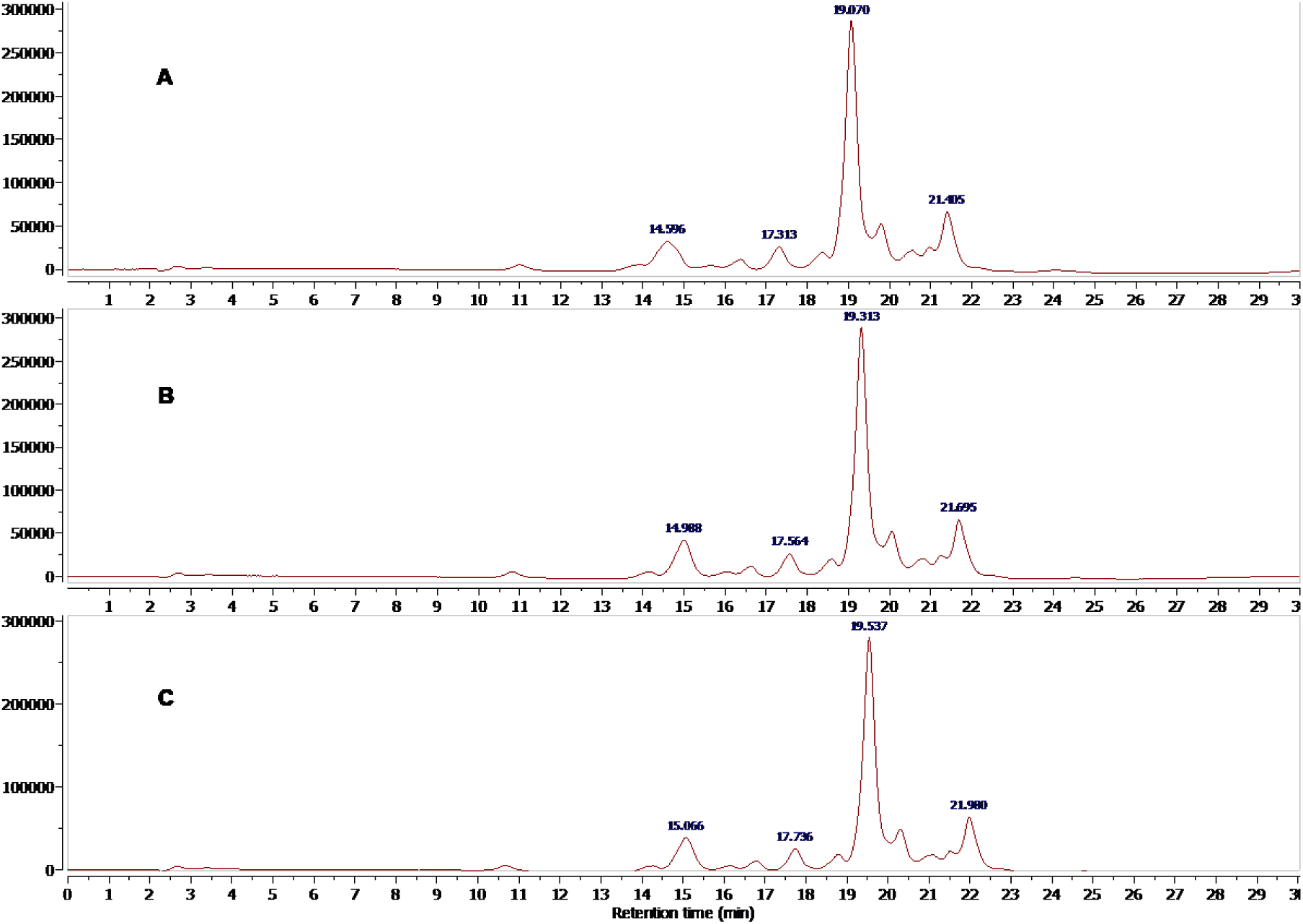
Effect of buffer concentration on QC sample. Column ZIC-cHILIC, 150 x 2.1 mm, 3 µm. (**A**) c(NH_4_Ac) = 20 mM. (**B**) c(NH_4_Ac) = 23 mM. (**C**) c(NH_4_Ac) = 25 mM.

**Figure 7.**
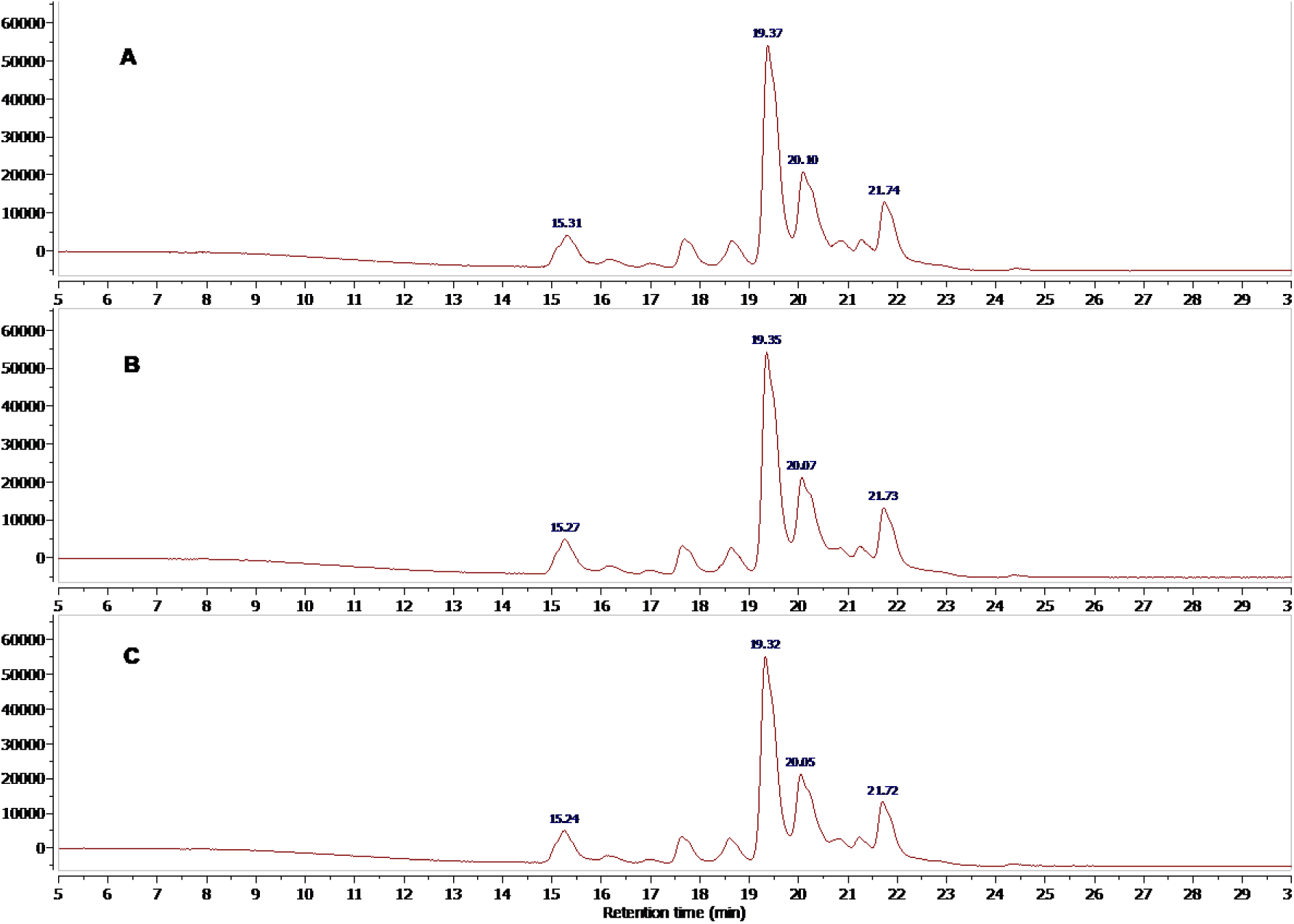
Effect of buffer pH on QC sample. Column ZIC-cHILIC, 150 x 2.1 mm, 3 µm. (**A**) pH 6.64. (**B**) pH 6.91; C) pH 7.11.

Small increases in both temperature or buffer concentration led to negligibly broader analyte regions, i.e. the first analyte (in our case IS) eluted earlier and the last analyte (in our case pppGpp) eluted later. In the case of temperature this phenomenon was constant for a wide temperature range [44] whereas for buffer concentration it was apparent only above certain concentration value. At lower concentration levels the retention factor decreased with increasing buffer concentration, whereas at higher concentration levels the retention factor increased with increasing buffer concentration [45] (compare **Fig. 3**).

Small pH changes (0.2 – 0.3 units) around neutral pH resulted in virtually no changes in analyte mobility as shown on **Fig. 8**.

**Figure 8.**
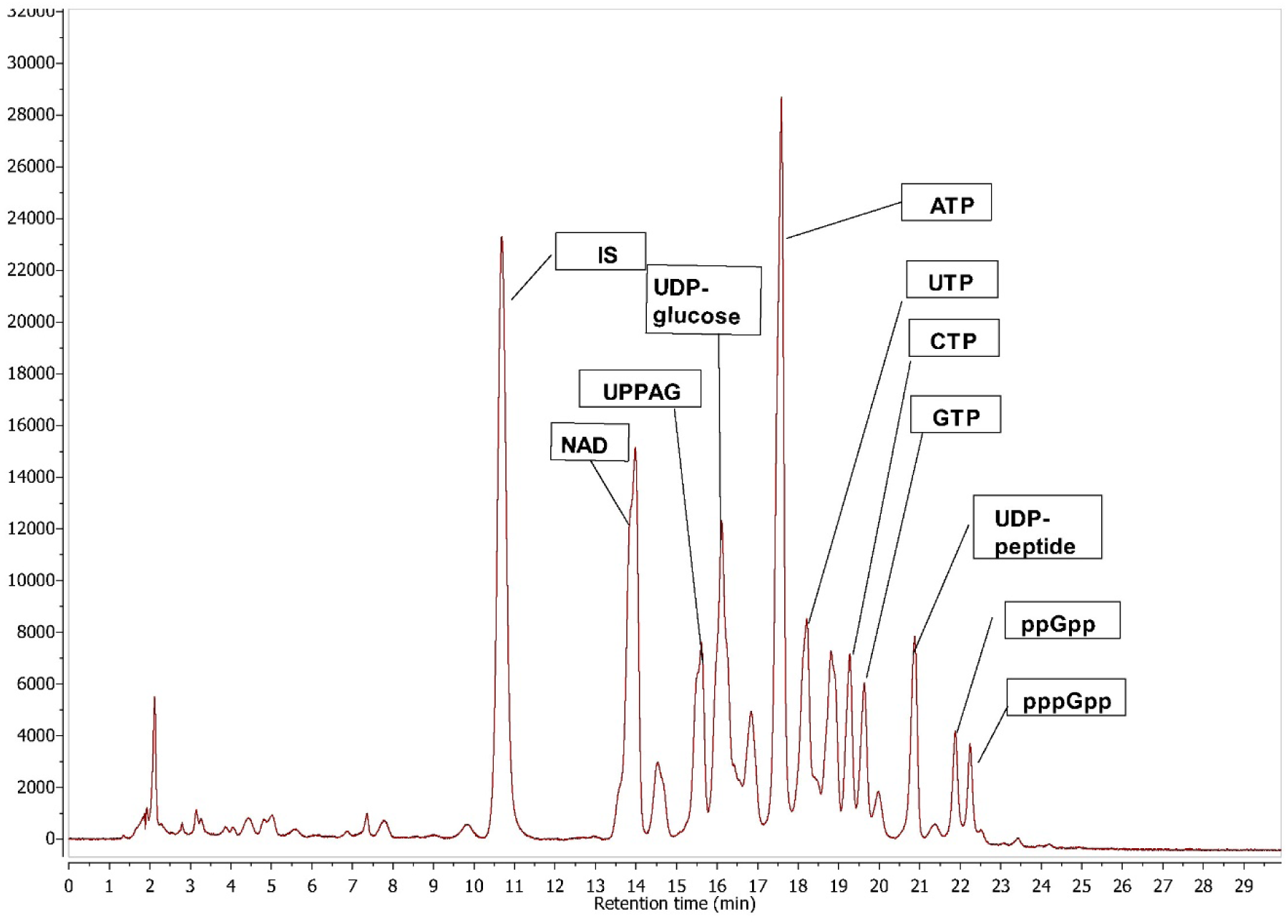
Chromatogram of the *E. coli* K12 extract after induction of acute amino acid starvation by 150 μg/ml mupirocin. Column ZIC-cHILIC, 150 x 2.1 mm, 3 µm. UDP-peptide = UDP-*N*-acetylmuramoyl-L-alanyl-D-glutamyl-6-carboxy-L-lysyl-D-alanyl-D-alanine, UPPAG = UDP-α-*N*-acetyl-D-glucosamine. Analytical parameters: see Materials and Methods section.

The composition of injection solvent and injection volume [46, 47] are known to affect separation in the HILIC mode. We tested different injection volumes and various percentages of acetonitrile (25%, 50% and 75%) with two column sizes (ZIC HILIC 150 mm, 250 mm) and two column types (ZIC-HILIC and ZIC-cHILIC). The results showed, that the smaller injection volume and the higher organic content, the better peak shape is obtained, as was expected. But another finding is connected to polarity of analyte, column dimension and concentration of buffer in mobile phase. The latter the compound eluted, the less affected peak shape is. Buffer content has the same effect, i.e. higher buffer content improved peak shape regardless of the injection solvent composition. More detailed discussion about this phenomenon is in *Injection solvent and volume section*.

### Matrix effects

Matrix effects (ME) are present in every complex sample, especially when it contains biological matrices [37]. Therefore, it is necessary to estimate the degree of ME on the obtained results. As the applied extraction method (1 M acetic acid) provided a clean extract without interfering peaks, we did not struggle with severe signal suppression. To prove this, we followed the recommended protocol [37] and evaluated ME on a standard mixture in 50% acetonitrile, blank growth medium and bacterial extract (see Materials and Methods). The difference between all three conditions in the estimated amount of IS was less then 5% and ME calculated according to *Material and methods* was calculated to 100. 5%, i.e. only a negligible signal enhancement was present. For ESI+ we determined ME as 89%, i.e. a moderate signal suppression had occurred. These results are in good agreement with the known characteristic of negative and positive ionization modes: negative ionization mode is known to cause signal suppression to a lower extent than positive ionization [48].

### Application of the method to bacterial extracts

As an indicator of the correctness of the obtained results, we used the Adenylate Energy Charge (AEC) ratio (Equation 1) [49], a key parameter that should remain about 0.8-0.9 for viable cells. Low AEC is a strong indicator of sample degradation.

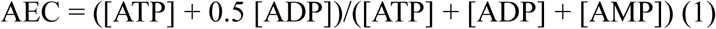

To evaluate the method on biological samples, we used cells of the well characterized *E. coli* K12 strain, using two physiological states: (i) non-stressed exponential phase cultures and (ii) mupirocin-challenged stringent cultures acutely starved for amino acid isoleucine and accumulating high levels of (p)ppGpp. The cells were filter-harvested and extracted as described in *Harvesting and extraction of bacterial cells*. Next, we determined the extraction efficiency using isotopically labeled ^13^C_10_-ATP as 64%. The apparent concentrations of analytes were then adjusted by this coefficient. This value was similar to Buckstein [20] who harvested the cells with formic acid and extracted nucleotides by vortexing followed by spinning and gel filtration, and who had estimated the overall recovery to be 71%. Our experimental results of nucleotide pools in exponentially growing *E. coli* in liquid LB media at OD_600_ 0.45 are presented in **Tab. 1** and plotted in **Fig. 8**. Comparison of intracellular concentration of published data with our results could be found in **Table S5** in the Supplementary material. It is evident, that there are substantial differences in absolute numbers amongst the individual reports, but relative distribution of particular nucleotides is significantly more similar (**Tab. S6**), with a few exceptions (e.g. the dTTP estimate by Bennett *et al.* [27]). The discrepancy in absolute numbers could be caused mainly by recalculation to volume of bacterial cell. For estimation of cell number it is possible to use flow cytometry [27] or plate counting [13]. In this report we used our previously published estimates for cell number and volume for *E. coli* exponentially growing in liquid LB media (see *Cell volume calculation* section and [13]).

**Table 1.** Intracellular concentration of 28 (deoxy)nucleotides in exponentially growing *E. coli* with (mupirocin +) and without (mupirocin -) induced stringent response. Concentration of mupirocin for induction of the stringent response was 150 µM. Based on the published data [13, 35] we approximated cell number for *E. coli* growing in LB media at the OD_600_ = 0.45 to 3.9 × 10^8^ and the cell volume to 1 × 10^-15^ l. AEC stands for Adenylate Energy Charge.

| compound | Intracellular concentration (μM) | |
| --- | --- | --- |
|  | Mupirocin - | Mupirocin + |
| ATP | 2024.5 | 1303.1 |
| ADP | 494.4 | 454.2 |
| AMP | 61.7 | 80.5 |
| GTP | 755.0 | 278.8 |
| GDP | 301.6 | 158.9 |
| GMP | 47.7 | 34.1 |
| cGMP | 2.5 | 4.4 |
| CTP | 520.2 | 340.6 |
| CDP | 411.7 | 526.4 |
| CMP | 117.0 | 172.2 |
| UTP | 242.7 | 162.0 |
| UDP | 230.2 | 244.1 |
| UMP | < LOQ | < LOQ |
| ITP | 871.1 | 495.8 |
| IDP | 126.6 | 111.4 |
| IMP | 115.6 | 21.1 |
| NAD | 698.0 | 1092.3 |
| NADH | 56.1 | 80.2 |
| NADP | < LOD | < LOD |
| NADPH | 126.9 | 109.5 |
| dATP | 121.7 | 78.9 |
| dAMP | 0.0 | 0.0 |
| dGTP | 88.3 | 28.0 |
| dGMP | < LOD | < LOD |
| dCTP | 111.3 | 61.6 |
| dCDP | 6.1 | 8.4 |
| dTTP | 553.4 | 988.1 |
| ppGpp | < LOQ | 793.8 |
| pppGpp | < LOQ | 459.8 |
| **AEC** | **0.88** | **0.83** |

## Discussion

We developed a reproducible and versatile method for determining nucleotide pools in bacterial cells using HPLC-single MS in the HILIC mode. To our best knowledge, this is the first application of this a methodology to analysis of bacterial nucleotide pools. The primary analytical setup consisted of HPLC system coupled with a single quadrupole mass spectrometer and PDA detector. The key aspects of the protocol, especially those requiring extensive optimization, are discussed below.

### Injection solvent and volume

Composition of injection solvent and injection volume are critical factors that could cause significant decrease of separation efficiency [46, 47, 50]. Good practice in RP chromatography is to inject sample in solvent close to the initial mobile phase composition. However, highly polar analytes, separated on HILIC, are poorly soluble in high acetonitrile content. For some analytes, mainly for those with adenine base, a double peak was observed when injected in pure water. Acetonitrile or salt addition suppressed this splitting. 50% of acetonitrile appeared to be sufficient to obtain well-defined peak shapes for all tested compounds. Even higher organic content did not improve either the peak shape or the recovery. These findings are in contrast to a study on unmodified Ethylene Bridged Hybrid (BEH) column [46] where relative strength of the injection solvent was reported crucial. The difference is likely due to differences in prior separation steps, stemming from the preferential role of the partitioning mechanism for unmodified BEH particles, and the contribution of week electrostatic interactions to ZIC-HILIC [51].

Furthermore, and importantly, total volume of injected water should be kept as low as possible. Higher injection volumes affect peak shapes at zwitterionic columns more than composition of the injection solvent does. This is in agreement with published studies [46, 47, 50]. Nevertheless, these studies do not mention that the degree to which the peak is affected depends also on the mobile phase composition and stationary phase chemistry. Peak shapes of well-retained compounds, (those eluted by up to 50% of water) are almost unaffected, regardless of the stationary phase used. However, peaks of early eluted compounds could broaden readily or show severe tailing.

Finally, we investigated the effect of injection solvent and its volume in dependence on the column size and chemistry. Our investigation on three columns (ZIC-HILIC 150 × 2.1 mm, 250 × 2.1 mm and ZIC-cHILIC 150 × 2.1 mm) revealed that the volume of injected water in respect to column void volume and concentration of the used buffer was the most crucial parameter. Higher void volume resulted in higher retention times and thus injection solvent affected less the peak shapes whereas higher buffer concentrations easily reduced the negative impact of added water. Dependencies of retention time on these parameters for ATP, ADP, AMP and pppGpp are shown in **Tab. 2.** RT of AMP strongly depended on injection solvent and higher amounts of water in the injection solvent often led to broad and split peaks. On the other hand, even at the ZIC-HILIC column (150 × 2.1 mm) the last eluted peak of pppGpp remained unaffected by solvent composition.

**Table 2.** Comparison of chromatographic behavior of selected analytes on different column chemistries and dimension. c (NH_4_Ac) concentration of ammonium acetate buffer in MF; V_0_ = column void volume, calculated based on retention time of unretained compound (toluene) and flow rate.

|  | Analyte | ZIC-HILIC,<br>250 x 2.1 mm | ZIC-HILIC,<br>150 x 2.1 mm | ZIC-cHILIC,<br>150 x 2.1 mm |
| --- | --- | --- | --- | --- |
| Reduced<br>retention<br>time<br>(min) | ATP | 14.05 | 11.72 | 16.94 |
|  | ADP | 12.07 | 11.04 | 15.27 |
|  | AMP | 8.76 | 8.11 | 12.88 |
|  | pppGpp | 20.01 | 14.61 | 21.33 |
| $c$ ($\text{NH}_4\text{Ac}$) (mM) | | 10 | 10 | 25 |
| $V_0$ (ml) | | 0.436 | 0.296 | 0.296 |

### Irreproducible retention times

Irreproducible retention times were mainly caused by insufficient column equilibration. We strongly recommend that before starting gradient elution with a new column or mobile phase composition, at least ten gradient full-time programs should be performed [50].

Another cause of irreproducible retention times between analyses could stem from different mobile phase composition as zwitterionic phase is highly sensitive to buffer pH and salt concentration. Moreover, HILIC is generally sensitive to water content and even a small variance in water volume can lead to unequal retention times mainly for early eluting compounds. This may cause problems if a binary gradient pump is used because in this case it is necessary to keep exact composition of the organic mobile phase. To maintain the same ionic strength during the entire separation process, and thus overcome corrupted peak shapes in the first part of the analysis, acetonitrile has to be diluted. This dilution has to reach at least 10% of water content for 10 mM acetate buffers, and 20% of water content for >20 mM acetate buffers. Ammonium acetate is insoluble in aprotic solvents and its insufficient solubility can lead to its precipitation during analysis and may damage the (U)HPLC system.

### On-column phosphate loss

During evaluation of the ^13^C_10_-ATP experiment, which addressed extraction recovery, we observed also the signal of ^13^C_10_-ADP at the intensity of one tenth of the intensity of ^13^C_10_-ATP. The cleavage of the γ-phosphate took place on the column and its extent depended on buffer concentration and type of the stationary phase.

As shown on **Fig. 9**, the higher salt concentration the stronger decomposition occurred. Triphosphate decomposition on the negatively charged surface of the ZIC-HILIC column was more concentration dependent than on the positively charged surface of ZIC-cHILIC. The γ-phosphate cleavage was not specific only for ATP but for other triphosphates and (p)ppGpp. The determined concentrations of diphosphates using the ZIC-cHILIC column were thus overestimated by 5-10% when compared to published data (**Tab. S5 and S6**).

**Figure 9.**
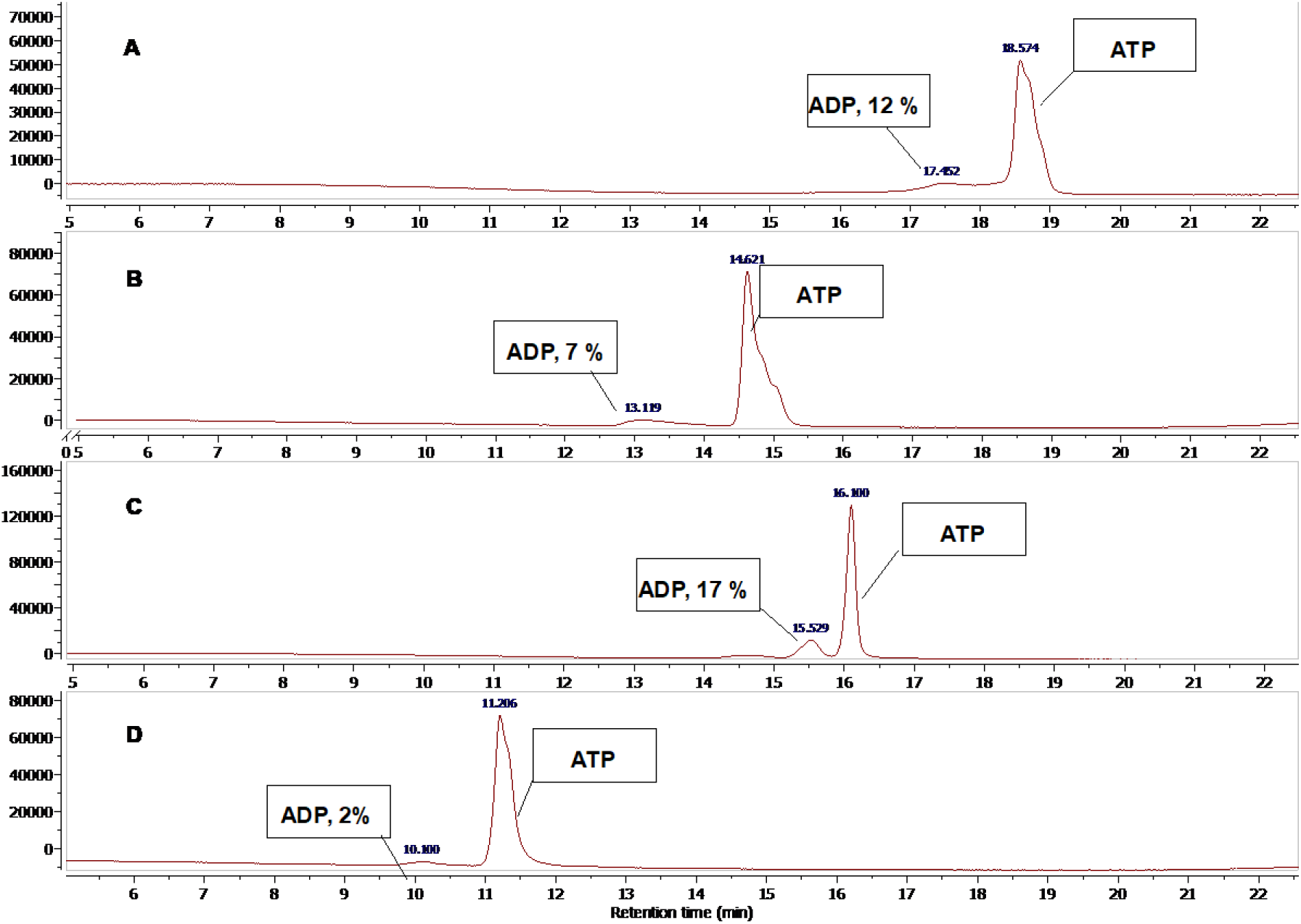
On column decomposition of adenosine triphosphate on two columns at two different buffer concentration. (**A**) ZIC –cHILIC column, 25 mM NH_4_Ac, gradient 1. (**B**) ZIC –cHILIC column, 10 mM NH_4_Ac, gradient 2. (**C**) ZIC –HILIC column, 25 mM NH_4_Ac, gradient 1. (**D**) ZIC –cHILIC column, 25 mM NH_4_Ac, gradient 2. All columns dimension: 150 x 2.1 mm, flow rate: 0,15 mL/min, injection volume: 2 µm. Gradient 1 = 0-2 min 20% B, 20-40% B in 17 min; Gradient 2= 0-3 min 100% B, 100-70% B in 20 min, hold till 25 min; flow rates = 150 µl/min. Column and MF specifications in Materials and Method section.

### Drifting MS signal intensity

Drifting MS signal intensity is a poorly understood phenomenon. Usually, the increase in the MS signal for the same compound and the same MS settings over a time period indicate that there is something wrong with the MS detector condition, such as room temperature, vacuum level *etc*. In some experiments with older column (during both standard and biological sample measurements) we observed continual increase in the MS response without any difference in internal or external conditions. As the MS signal of IS increased as well, the corrected response remained constant. Washing columns with pure solvents without buffer could decrease the signal drifting for a couple of injections, but on the other hand, it could also cause asymmetrical peaks since the active sites of the column were not saturated yet. Replacing an older column with a new one solves this issue. Why the MS signal is drifting for older columns over the time is not clear yet but using a well selected internal standard eliminates incorrect results.

### Calibration curve

The calibration curve is usually calculated with a linear regression model, which provides the best fits for the majority of obtained data, especially for those from a UV detector. However, the MS signal does not exhibit linear behavior, especially at higher concentrations close to the detector saturation limit or when adducts are formed in dependence on analyte concentration. There are two other most common possibilities how to interpolate the calibration points – quadratic function or linear log-log function. In our case, the former is suitable for triphosphates or polyphosphates ionized in ESI- (**Fig. 10**) whereas the latter for monophosphates ionized in ESI+.

**Figure 10:**
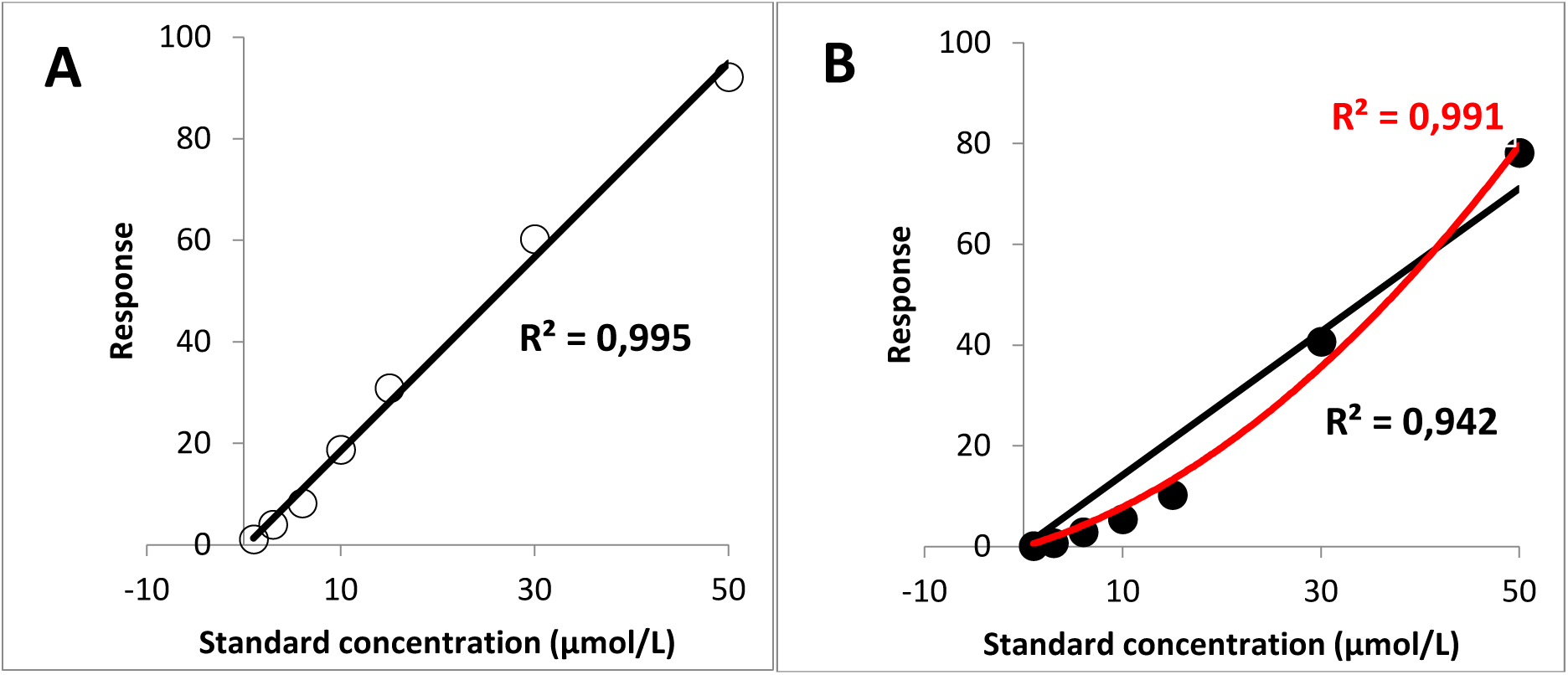
Regression model for calibration curve with appropriate coefficient of determination. (**A**) Linear regression for AMP; (**B**) Linear (black line) and quadratic (red line) regression for ATP.

### Conclusions

We have developed and validated a simple, fast and versatile method for determination of nucleotide concentrations in bacterial cell samples using a zwitterionic column with single MS detection. The method relies on a positively charged ZIC-cHILIC column that proved to be most efficient for complex nucleotide screening and was able to distinguish even between stringent response alarmones ppGpp and pppGpp. A major advantage of this method is the use of single MS detection that is currently a standard equipment in many laboratories. It allows for precise identification of analytes of interest without special knowledge about the instrumentation. The usage of broad range of isotopically labeled standards, as is common is metabolomic analysis, is not necessary here – a single IS is sufficient. Finally, the sensitivity of this methods is higher compared to methods using only simple UV detection. Due to its simplicity and versatility, this ZIC-cHILIC-MS method promises to be applicable to screening nucleotide pools in various bacterial species and be of broad interest to researchers studying cellular metabolism.

## Supporting information

Article

## List of abbreviations

(c-di-AMP): cyclic diadenylate
(c-di-GMP): cyclic diguanylate
ApxN: dinucleoside polyphosphates
(pppGpp): guanosine pentaphosphate
(ppGpp): guanosine tetraphosphate
(LB): lysogeny broth
(SR): stringent response
(AEC): Adenylate Energy Charge

## Acknowledgements

This work was supported by Czech ministry of health (Grant No.17-29680A to D.R. and L.K.) Czech Science Foundation grant number (grant No. 15-11711S to D.R.), EU and Czech ministry of Education and Sport via JPIAMR (grant No. 8F19006 to D.R. and V.H.), Swedish Research council (grant 2017-03783 to VH) and Charles University, Department of Analytical Chemistry – Specific University Research No. SVV260440 to E.Z.

