## Supplementary material for "Analysis of Nucleotide Pools in Bacteria Using HPLC-MS in HILIC Mode": Article

### Supplementary data

**Table S1:** QC mixture composition and measured intermediate precision with standard deviation (n=3). Column ZIC-cHILIC, 150 x 2.1 mm.

| Compound | In vitro concentration (µmol/L) | Measured concentration (µmol/L) | bias (%) |
|---|---|---|---|
| **ATP** | 150 | 143 | 8% |
| **ADP** | 25 | 45 | 13% |
| **AMP** | 10 | 26 | 5% |
| **dATP** | 17.5 | 22 | 6% |
| **GTP** | 60 | 53.5 | 1% |
| **GDP** | 10 | 7.5 | 9% |
| **GMP** | 0.2 | 0.5 | 16% |
| **dGTP** | 12 | 13.5 | 5% |
| **CTP** | 50 | 44.5 | 2% |
| **UTP** | 50 | 54.5 | 1% |
| **UDP** | 9 | 11 | 0% |
| **NAD** | 30 | 28 | 5% |
| **NADH** | 15 | 8.4 | 20% |
| **NADP** | 10 | 11.6 | 24% |
| **NADPH** | 10 | 6.9 | 27% |
| **ppGpp** | 3 | 3.5 | 35% |
| **pppGpp** | 2 | 3.0 | 36% |

**Table S2**: Validation parameters for ZIC-HILIC column. (250 x 2.1 mm, 3.5 µm; flow rate = 0.15 mL/min). LOD = Limit of detection, LOQ = Limit of quantification, N.D. = Not determined.

| Analyte | $[M-H]^-$ | LOD (µmol/L) | LOQ (µmol/L) | Tf | As | r.t. (min) |
|---|---|---|---|---|---|---|
| cGMP | **344.05** | <1 | 1 | 1.17 | 1.35 | 7.65 |
| NADH | **644.11** | 2 | 5 | 1.11 | 1.21 | 8.33 |
| AMP | **346.05** | 1 | 2 | 0.81 | 0.63 | 10.92 |
| dGMP | **346.06** | 1 | 2 | 1.15 | 1.28 | 11.30 |
| NAD | **662.09** | 2 | 5 | 1.26 | 1.55 | 12.51 |
| UMP | **323.04** | 1 | 2 | 0.84 | 0.68 | 13.25 |
| IMP | **347.04** | 2 | 3 | 1.12 | 1.23 | 13.93 |
| ADP | **426.02** | 3 | 5 | 1.17 | 1.32 | 13.94 |
| dTTP | **480.98** | 3 | 5 | 1.00 | 1.01 | 14.68 |
| dATP | **489.99** | 5 | 10 | 1.11 | 1.20 | 14.98 |
| UDP | **402.99** | 2 | 3 | 1.06 | 1.11 | 15.33 |
| NADPH | **744.08** | 3 | 10 | 1.02 | 1.04 | 15.53 |
| ATP | **505.99** | 3 | 5 | 1.37 | 0.59 | 15.82 |
| CMP | **322.05** | 1 | 2 | 1.12 | 1.41 | 16.30 |
| IDP | **427.00** | 5 | 10 | 0.95 | 0.89 | 16.48 |
| GMP | **362.05** | 1 | 2 | 1.17 | 1.34 | 16.61 |
| dITP | **490.97** | 3 | 5 | 1.33 | 1.67 | 16.64 |
| UTP | **482.97** | 3 | 5 | 1.78 | 2.58 | 17.01 |
| CDP | **402.01** | 3 | 5 | 0.96 | 0.92 | 17.30 |
| ITP | **506.99** | 5 | 10 | 1.26 | 1.52 | 17.50 |
| NADP | **742.06** | 3 | 10 | 1.28 | 1.52 | 18.11 |
| GDP | **442.02** | 3 | 5 | 1.14 | 1.58 | 18.25 |
| dGTP | **505.99** | 5 | 10 | 1.30 | 1.60 | 18.30 |
| CTP | **481.97** | 3 | 5 | 1.43 | 1.85 | 18.83 |
| pGp | **426.02** | 5 | 10 | 1.47 | 1.95 | 19.46 |
| GTP | **521.98** | 5 | 10 | 1.73 | 2.38 | 19.47 |
| pGpp | **521.98** | 5 | 10 | 1.06 | 1.11 | 21.25 |
| ppGpp | **601.97** | 10 | 30 | 1.28 | 1.59 | 21.80 |
| pppGpp | **681.97** | 10 | 30 | 1.26 | 1.48 | 22.05 |
| IS250 | **249.11** | N.D. | N.D. | 1.12 | 1.22 | 22.65 |

**Table S3:** Validation parameters for ZIC-cHILIC column (150 x 2.1 mm, 3 µm; flow rate 0.15 mL/min). LOD = Limit of detection, LOQ = Limit of quantification, N.D. = Not determined.

| Analyte | $[M-H]^-$ | LOD (µmol/L) | LOQ (µmol/L) | Tf | As | r.t. (min) |
|---|---|---|---|---|---|---|
| **cGMP** | **344.05** | <0.2 | 0.2 | 0.80 | 0.61 | *9.61* |
| **IS250** | **249.11** | N.D. | N.D. | 1.18 | 1.35 | *11.35* |
| **NADH** | **644.11** | 0.6 | 1 | 1.18 | 1.36 | *13.79* |
| **AMP** | **346.05** | <0.2 | 0.2 | 1.08 | 1.17 | *14.82* |
| **NAD** | **662.09** | 0.2 | 0.6 | 0.97 | 0.93 | *14.97* |
| **UMP** | **323.04** | 0.2 | 0.6 | 0.97 | 0.91 | *15.77* |
| **IMP** | **347.04** | 0.2 | 0.6 | 0.97 | 0.93 | *16.50* |
| **dGMP** | **346.06** | 0.2 | 1 | 0.90 | 0.79 | *16.75* |
| **ADP** | **426.02** | 0.2 | 0.6 | 1.06 | 1.12 | *17.13* |
| **dTTP** | **480.98** | 0.6 | 2 | 0.98 | 0.90 | 17.44 |
| **CMP** | **322.05** | <0.2 | 0.2 | 0.96 | 0.93 | *17.68* |
| **UDP** | **402.99** | 0.6 | 2 | 1.14 | 1.29 | *17.82* |
| **NADPH** | **744.08** | 2.5 | 5 | 0.96 | 0.94 | *17.89* |
| **GMP** | **362.05** | <0.2 | 0.2 | 1.05 | 1.11 | *17.92* |
| **ITP** | **506.99** | 0.6 | 2 | 1.18 | 1.36 | *18.02* |
| **dATP** | **489.99** | 0.2 | 1 | 1.05 | 1.08 | *18.11* |
| **IDP** | **427.00** | 0.6 | 2 | 0.96 | 0.93 | *18.37* |
| **ATP** | **505.99** | 0.2 | 1 | 1.20 | 1.38 | *18.74* |
| **CDP** | **402.01** | 0.2 | 3 | 1.13 | 1.24 | *19.09* |
| **dITP** | **490.97** | 0.2 | 1 | 1.15 | 1.30 | *19.15* |
| **UTP** | **482.97** | 0.6 | 2 | 1.03 | 1.05 | *19.32* |
| **GDP** | **442.02** | 0.2 | 1 | 0.98 | 0.97 | *19.46* |
| **dGTP** | **505.99** | 0.6 | 2 | 1.00 | 1.00 | *20.05* |
| **NADP** | **742.06** | 0.6 | 1 | 1.01 | 1.02 | *20.39* |
| **CTP** | **481.97** | 0.6 | 3 | 1.13 | 1.25 | *20.40* |
| **GTP** | **521.98** | 0.6 | 2 | 0.98 | 0.98 | *20.78* |
| **pGp** | **442.02** | 0.2 | 1 | 1.21 | 1.42 | *21.13* |
| **pGpp** | **521.98** | 0.6 | 2 | 1.00 | 0.99 | *21.95* |
| **ppGpp** | **601.97** | 2 | 5 | 1.07 | 1.13 | *22.66* |
| **pppGpp** | **681.97** | 2 | 5 | 1.32 | 1.66 | *23.37* |

**Table S4:** Retention factor ($k_A$), separation factor (α) and resolution ($R_s$) of critical pair of analytes.

| Analyte | $[M-H]^-$ | r.t. (min) | $k_A$ | α | Rs |
|---|---|---|---|---|---|
| **ATP** | **505.99** | *18.74* | 8.51 | 1.08 | 2.18 |
| **dGTP** | **505.99** | *20.05* | 9.18 | | |
| **AMP** | **346.05** | *14.82* | 6.52 | 1.15 | 1.74 |
| **dGMP** | **346.06** | *16.75* | 7.50 | | |
| **dATP** | **489.99** | *18.11* | 8.19 | 1.06 | 1.53 |
| **dITP** | **490.97** | *19.15* | 8.72 | | |
| **CTP** | **481.97** | *20.4* | 9.36 | 1.06 | 1.79 |
| **UTP** | **482.97** | *19.32* | 8.81 | | |
| **pppGpp** | **681.97** | *23.37* | 10.86 | 1.09 | 1.62 |
| **ppGpp** | **601.97** | *21.66* | 9.99 | | |
| **GTP** | **521.98** | *20.78* | 9.55 | 1.06 | 2.55 |
| **pGpp** | **521.98** | *21.95* | 10.14 | | |
| **GDP** | **442.017** | *19.46* | 8.88 | 1.10 | 1.99 |
| **pGp** | **442.017** | *21.13* | 9.73 | | |
| **NADH** | **664.11** | *13.79* | 6.00 | 1.10 | 1.38 |
| **NAD** | **662.09** | *14.97* | 6.60 | | |
| **NADPH** | **744.08** | *17.89* | 8.08 | 1.16 | 1.10 |
| **NADP** | **742.06** | *20.39* | 9.35 | | |

**Table S5:** Comparison of intracellular concentration of published data with our results.

| compound | Intracellular concentration (µmol/L) | | | | | |
|---|---|---|---|---|---|---|
| | ***S. enterica*, Bochner, 1982** | ***E.coli*, Buckstein, 2007** | ***E. coli*, Varik, 2017** | ***E. coli*, Bennett, 2009** | ***E. Coli*, No mupirocine** | **E. Coli, Mupirocine (150 ug/mL)** |
| **ATP** | 3000 | 3560 | 2200 | 9600 | 2024.5 | 1303.1 |
| **ADP** | 250 | 116 | 430 | 560 | 494.4 | 454.2 |
| **AMP** | 100 | | 88 | 280 | 61.7 | 80.5 |
| **dATP** | 175 | 181 | | 16 | 121.7 | 78.9 |
| **dAMP** | | | | 9 | 0.0 | 0.0 |
| **GTP** | 900 | 1660 | 900 | 4900 | 755.0 | 278.8 |
| **GDP** | 125 | 203 | 160 | 680 | 301.6 | 158.9 |
| **GMP** | 20 | | 58 | 24 | 47.7 | 34.1 |
| **dGTP** | 120 | 92 | | | 88.3 | 28.0 |
| **dGMP** | | | | 51 | < LOD | < LOD |
| **IMP** | 160 | | | 270 | 115.6 | 21.1 |
| **CTP** | 500 | 325 | 570 | 2700 | 520.2 | 340.6 |
| **CDP** | 80 | | | | 411.7 | 526.4 |
| **CMP** | | | | 360 | 117.0 | 172.2 |
| **dCTP** | 60 | 184 | | 35 | 111.3 | 61.6 |
| **UTP** | 900 | 667 | 970 | 8300 | 242.7 | 162.0 |
| **UDP** | 90 | 54 | | 1800 | 230.2 | 244.1 |
| **UMP** | 140 | | | | < LOQ | < LOQ |
| **dTTP** | 80 | 256 | | 4600 | 553.4 | 988.1 |
| **NAD** | 300 | | | 2600 | 698.0 | 1092.3 |
| **NADH** | | | | 83 | 56.1 | 80.2 |
| **NADP** | 200 | | | 2 | < LOD | < LOD |
| **NADPH** | | | | 12 | 126.9 | 109.5 |
| **ppGpp** | 30 | | 120 | | < LOQ | 793.8 |
| **pppGpp** | 20 | | | | < LOQ | 459.8 |
| **ITP** | | | | | 871.1 | 495.8 |
| **IDP** | | | | | 126.6 | 111.4 |
| **cGMP** | | | | | 2.5 | 4.4 |
| **dCDP** | | | | | 6.1 | 8.4 |
| **AEC** | **0.93** | **0.98** | **0.88** | **0.95** | **0.88** | **0.83** |

**Table S6:** Comparison of relative intracellular concentration of published data with our results; ATP = 100 %.

| compound | Intracellular concentration (µmol/L) | | | | | |
|---|---|---|---|---|---|---|
| | ***S. enterica*, Bochner, 1982[1]** | ***E.coli*, Buckstein, 2007[2]** | ***E. coli*, Varik, 2017[3]** | ***E. coli*, Bennett, 2009[4]** | ***E. Coli*, No mupirocine** | ***E. Coli*, Mupirocine (150 ug/mL)** |
| **ATP** | 100.0% | 100.0% | 100.0% | 100.0% | 100.0% | 100.0% |
| **ADP** | 8.3% | 3.3% | 19.5% | 5.8% | 24.4% | 34.9% |
| **AMP** | 3.3% | | 4.0% | 2.9% | 3.0% | 6.2% |
| **dATP** | 5.8% | 5.1% | | 0.2% | 6.0% | 6.1% |
| **dAMP** | | | | 0.1% | | |
| **GTP** | 30.0% | 46.6% | 40.9% | 51.0% | 37.3% | 21.4% |
| **GDP** | 4.2% | 5.7% | 7.3% | 7.1% | 14.9% | 12.2% |
| **GMP** | 0.7% | | 2.6% | 0.3% | 2.4% | 2.6% |
| **dGTP** | 4.0% | 2.6% | | | 4.4% | 2.1% |
| **dGMP** | | | | 0.5% | 0.0% | 0.0% |
| **IMP** | 5.3% | | | 2.8% | 5.7% | 1.6% |
| **CTP** | 16.7% | 9.1% | 25.9% | 28.1% | 25.7% | 26.1% |
| **CDP** | 2.7% | | | | 20.3% | 40.4% |
| **CMP** | | | | 3.8% | 5.8% | 13.2% |
| **dCTP** | 2.0% | 5.2% | | 0.4% | | |
| **UTP** | 30.0% | 18.7% | 44.1% | 86.5% | 12.0% | 12.4% |
| **UDP** | 3.0% | 1.5% | | 18.8% | 11.4% | 18.7% |
| **UMP** | 4.7% | | | | | |
| **dTTP** | 2.7% | 7.2% | | 47.9% | 27.3% | 75.8% |
| **NAD** | 10.0% | | | 27.1% | 34.5% | 83.8% |
| **NADH** | | | | 0.9% | 2.8% | 6.2% |
| **NADP** | 6.7% | | | 0.02% | | |
| **NADPH** | | | | 0.1% | 6.3% | 8.4% |
| **ppGpp** | 1.0% | | 5.5% | | | 60.9% |
| **pppGpp** | 0.7% | | | | | 35.3% |
| **ITP** | | | | | 43.0% | 38.0% |
| **IDP** | | | | | 6.3% | 8.5% |

[1] B.R. Bochner, B.N. Ames, Complete Analysis of Cellular Nucleotides by Two-Dimensional Thin-Layer Chromatography, Journal of Biological Chemistry 257(16) (1982) 9759-9769.
[2] M.H. Buckstein, J. He, H. Rubin, Characterization of nucleotide pools as a function of physiological state in Escherzchia coli, Journal of Bacteriology 190(2) (2008) 718-726.
[3] V. Varik, S.R.A. Oliveira, V. Hauryliuk, T. Tenson, HPLC-based quantification of bacterial housekeeping nucleotides and alarmone messengers ppGpp and pppGpp, Scientific Reports 7 (2017).
[4] B.D. Bennett, E.H. Kimball, M. Gao, R. Osterhout, S.J. Van Dien, J.D. Rabinowitz, Absolute metabolite concentrations and implied enzyme active site occupancy in Escherichia coli, Nature Chemical Biology 5(8) (2009) 593-599.
